# TreeProfiler: Large-scale metadata profiling along gene and species trees

**DOI:** 10.1101/2023.09.21.558621

**Authors:** Ziqi Deng, Ana Hernández-Plaza, Adrián A. Davín, Jaime Huerta-Cepas

## Abstract

Profiling biological traits along gene or species tree topologies is a well-established approach in comparative genomics, widely employed to infer gene function from co-evolutionary patterns (phylogenetic profiling), reconstruct ancestral states, and uncover ecological associations. However, existing profiling tools are typically tailored to specific use cases, have limited scalability for large datasets, and lack robust methods to aggregate or summarize traits at internal tree nodes. Here, we present TreeProfiler, a tool for automated annotation and interactive exploration of hundreds of features along large gene and species trees, with seamless summarization of mapped traits at internal nodes. TreeProfiler supports the profiling of custom continuous and discrete traits, as well as ancestral character reconstruction and phylogenetic signal tests. It also integrates commonly used genomic features, including multiple sequence alignments, protein domain architectures, and functional annotations. We demonstrate TreeProfiler’s utility beyond traditional phylogenetic profiling by analyzing functional diversification and domain architectures within the pyruvate flavodoxin/ferredoxin oxidoreductase gene family (13,297 genes), and by profiling the relative abundance of 124,295 bacterial and archaeal species across 51 biomes. TreeProfiler is open-source and freely available at https://github.com/compgenomicslab/TreeProfiler.

## INTRODUCTION

Phylogenetic tree profiling is a comparative genomics approach used to analyze the distribution of one or more traits across species or genes while taking their evolutionary relationships into account. Initially proposed to infer functional associations between proteins based on their co-evolutionary patterns (Pellegrini et al. 1999), phylogenetic profiling has become a versatile exploration tool with a wide range of applications (Moi and Dessimoz 2023), from inferring protein interactions and identifying synapomorphic traits (Peng et al. 2023) to detecting horizontal transfer (Ravenhall et al. 2015) and convergent evolution events (Rey et al. 2018). In microbial ecology, phylogenetic profiling has also become useful for studying functional and ecological patterns of genes and species across environmental samples (Asnicar et al. 2015).

Although this broad utility has led to the development of numerous bioinformatics tools, phylogenetic tree profiling remains challenging when applied to the large and complex data generated by current genomic studies. First, annotating phylogenetic trees often requires *ad hoc* pipelines and is typically restricted to small and medium datasets. In addition, most phylogenetic profiling tools tend to focus on traditional use cases, such as profiling orthologous groups (Hernández-Plaza et al. 2022; Rossier et al. 2024; Szklarczyk et al. 2025; Tran and Ebersberger 2025), protein function (Fang et al. 2021) or domain architectures (Cromar et al. 2016), offering limited options to profile custom metadata. Tree visualization software, including programming libraries such as ETE Toolkit and ggtree (Huerta-Cepas et al. 2016; Yu et al. 2017), standalone applications (Asnicar et al. 2015; Cantrell et al. 2021) and online platforms like iTOL or tvBOT, (Xie et al. 2023; Letunic and Bork 2024), are therefore the default option for exploring large and customly annotated phylogenetic trees. However, visual representation and interactive exploration of very large annotated phylogenies is a major challenge, as trees and metadata are usually too large to be rendered and iteratively navigated. Moreover, tree visualization software treats leaf-node features as static graphical elements, which can lead to misleading representations when plotting collapsed versions of large phylogenies (Supplementary Figure 1).

To address these limitations, we have developed TreeProfiler, a bioinformatics tool designed to automate the annotation of very large phylogenetic trees with custom features, facilitating its interactive visualization. TreeProfiler supports a wide range of metadata to be annotated and propagated across internal nodes of a phylogenetic tree, representing either discrete or continuous traits. Furthermore, it can easily incorporate functional annotations, taxonomic data, multiple sequence alignments, and protein domain architectures into the profiling options.

Finally, TreeProfiler provides seamless integration with methods for ancestral character reconstruction of discrete characters (Ishikawa et al. 2019; Revell 2024), phylogenetic signal tests (Ribeiro et al. 2023), and estimation of lineage-specific traits (Mendler et al. 2019). Interactive exploration of phylogenetic profiles is powered by ETE Toolkit v4.0, enabling efficient and interactive exploration of trees with hundreds of thousands of nodes.

## IMPLEMENTATION AND MAIN FEATURES

TreeProfiler is implemented as a command-line tool for annotating and visualizing phylogenetic trees with metadata. It provides two subcommands: *treeprofiler-annotate* for computing profiles and annotating metadata onto trees; and *treeprofiler-plot* for interactive visualization and exploration of profiled data.

### Computing phylogenetic profiles

The *treeprofiler-annotate* command is responsible for annotating each of the nodes in a phylogenetic tree (in Newick format), with custom metadata provided by the user, encoded as one or more data tables (in TSV format). Each row in the metadata tables is expected to contain the list of annotations of a particular terminal or internal tree node, while each column represents a different annotation track. Unless explicitly specified, TreeProfiler will detect the most suitable data type for each annotation track (columns). After initial annotation of the provided metadata, *treeprofiler-annotate* will perform a general scan of the tree topology and infer missing annotations for all internal nodes of the phylogeny, summarizing values under each internal branch according to the data type of each annotation track. For instance, continuous data types will be summarized using descriptive statistics, while discrete values will be used to calculate frequency distributions. For convenience, multi-value categorical annotations can be automatically converted into presence/absence matrices. TreeProfiler can also perform ancestral trait analysis on selected annotation tracks, including ancestral character reconstruction (ACR) by maximum parsimony or maximum likelihood (using PastML (Ishikawa et al. 2019)), phylogenetic signal tests using delta statistic (Ribeiro et al. 2023), and lineage specificity metrics for boolean traits. These methods can be used for one or more annotation tracks, resulting in the annotation of internal tree nodes with the corresponding values.

Special data types are supported to link common genomic information with tree topology, such as multiple sequence alignments, domain architectures, functional annotations and taxonomic assignments. Thus, TreeProfiler can automatically interpret taxonomic annotations based on NCBI Taxonomy IDs (Sayers et al. 2018), GTDB (Parks et al. 2021) and mOTUs (Dmitrijeva et al. 2025), which are further used to infer full lineage tracks, species names, taxonomic rank and last common ancestry along all tree nodes. Similarly, functional annotations and protein domain coordinates generated by eggNOG-mapper (Cantalapiedra et al. 2021), a widely used tool for functional annotation of genomes and metagenomes, are seamlessly integrated, allowing users to explore gene phylogenies alongside the distribution of, for example, Gene Ontology terms, KEGG annotations or Pfam protein domain architectures (Mistry et al. 2020). Additionally, multiple sequence alignment can be associated with their respective phylogenies, enabling automatic calculation of consensus sequences at internal nodes.

The output of *treeprofiler-annotate* is an annotated tree file with embedded metadata, which can be further used for programmatic analysis or custom visualization.

### Visualizing phylogenetic profiles and metadata

The *treeprofiler-plot* command allows users to interactively visualize and explore previously annotated trees, letting them decide and change the appropriate graphical layout used for each annotation track. The graphical user interface of *treeprofiler-plot* is based on the newest ETE Toolkit drawing engine (v4), which provides support for the interactive navigation of very large phylogenies (e.g., >100,000 tree nodes), as well as dynamic control of what annotation tracks are visible (Figure 1A).

**Figure 1.**
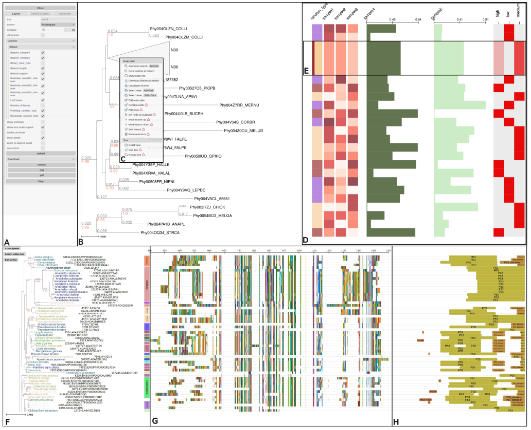
Overview of the *treeprofiler-plot* interface. **A)** Control panel for customizing visualization layouts, enabling annotation tracks, and managing search options. **B)** Phylogenetic tree viewer displaying support values (red) and branch lengths (gray). **C)** Node context menu with options for collapsing, pruning, rooting, and other per-node actions. **D**) Phylogenetic profiles of various data types visualized across the tree, including categorical traits (first column of color-coded rectangles), numeric traits displayed as heatmap gradients (columns 2–4) and bar plots (columns 5–6), as well as binary presence/absence profiles for categorical traits (final column). Column headers indicate the annotation track names. **E)** Summarized annotations for a collapsed clade: frequency distribution of categorical value is shown as stacked bar (column 1), averaged heatmap values (columns 2-4), averaged barplot values (columns 5–6) and presence/absence matrix gradients (positive-to-total ratio) at the last column. **F)** A phylogenetic tree annotated with taxonomic annotation and visualized with phylum-level classification using vertical color bands on corresponding clades. Leaf nodes are labeled with scientific names based on NCBI taxonomy. **G)** Multiple sequence alignment used to infer the tree. **H)** Most frequent protein domain architectures identified under each tree branch.

Continuous traits can be visualized as text labels, heatmaps, bar plots, or color gradients; while categorical traits can be shown as colored labels, symbols, or converted into heatmap-based presence/absence matrices (Figure 1D). Importantly, when clades are collapsed and leaf nodes are not visible, TreeProfiler will display the annotations inferred for internal nodes, representing the statistical summary of all values collapsed under each node. For instance, instead of graphically overlapping the representation of multiple leaf values, the average, maximum, minimum or frequency distribution of the grouped leaves will be shown (Figure 1E and Supplementary Figure 1). Special annotation tracks have their own dedicated layouts and graphical style configurations. Thus, taxonomic classifications, including common ancestry, are represented as vertical color bands aligned with their corresponding clades (Figure 1F); multiple sequence alignments and consensus sequence representations are dynamically adjusted to zoom level (Figure 1G); and the most frequent protein domain architecture is shown of each branch (Figure 1H).

Furthermore, *treeprofiler-plot* offers options for setting up custom node collapsing and highlighting rules based on tree node annotations. For example, large trees can automatically collapse at a specific taxonomic rank or at internal nodes that meet particular size or annotation value criteria.

Finally, to facilitate tree profile visualization of large data tables, the helper command *treeprofiler-desktop* is available, which will launch a web-based interface to guide the exploration of all annotation tracks available in a tree, as well as their possible visualization options and layouts offered by TreeProfiler.

## SCALABILITY & USE CASES

TreeProfiler is a versatile tool built for the daily exploration of phylogenetic trees and their associated metadata, regardless of their size or complexity. While small datasets can be handled with ease, TreeProfiler is particularly well suited for larger, more intricate data, making it ideal for two primary applications: (i) Gene tree-based phylogenetic profiling, enabling the exploration of subfunctionalization patterns, orthologous groups, and the distribution of custom genomic features; and (ii) Species tree-based profiling, allowing for the interactive exploration of functional, ecological, and evolutionary patterns along species phylogenies.

To demonstrate the first use case, we employed TreeProfiler to analyse the functional annotations and domain architectures of the *pyruvate flavodoxin/ferredoxin oxidoreductase* protein family (POR_N domain, PF01855). This large protein family is found in all domains of life and comprises various orthologous groups and known subfunctionalization events. We therefore collected 13,297 protein sequences containing the POR_N domain from the eggNOG v6 database (Hernández-Plaza et al. 2022), and inferred a multiple sequence alignment and phylogenetic tree using MAFFT v7.525 (Katoh and Standley 2013) and FastTree v2.1.11 (Price et al. 2010), respectively. We also annotated the 13,297 protein sequences, spanning 6,014 taxa, with the functional terms and PFAM domain coordinates inferred by eggNOG-mapper (Cantalapiedra et al. 2021). We then used *treeprofiler-annotate* to link the resulting tree with the original multiple sequence alignment, taxonomic information and eggNOG-mapper results. Interactive exploration of the annotated tree by *treeprofiler-plot* revealed various speciation and duplication events (Figure 2A) coinciding with putative subfunctionalizations based on the KEGG KO functional terms distribution across tree topology (Figure 2B) and the distinct pattern of domain architectures associated to each subgroup (Figure 2C).

**Figure 2.**
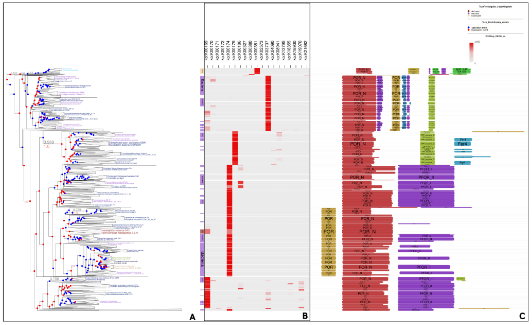
Phylogenetic profiling of the POR_N protein domain family with TreeProfiler. **A)** Phylogenetic tree of 13,297 POR_N containing protein sequences. Evolutionary events in the internal node indicate speciations (blue) or duplications (red). **B)** Heatmap showing orthology and functional assignments based on KEGG terms inferred by eggNOG-mapper. Color gradients represent the proportion of proteins annotated to each KO over the total number of descendant sequences within each collapsed clade. **C)** Dominant Pfam domain architectures under each node.

The second use case (species tree-based profiling) is illustrated by the use of TreeProfiler in the context of microbial ecology research. To demonstrate its scalability, we inferred, annotated and visualized the complete mOTUs reference taxonomy tree—currently comprising 124,295 archaeal and bacterial species (Dmitrijeva et al. 2025)—together with the abundance estimate of each reference species across 112,121 metagenomic samples from 51 biomes. To obtain the tree, we concatenated the alignments of the ten single-copy phylogenetic marker genes used by mOTUs across all the 124,295 species. Each marker was independently aligned using MAFFT (Katoh and Standley 2013) and trimmed with TrimAl (Capella-Gutiérrez et al. 2009) (-gt 0.1), resulting in a final concatenated alignment of 5,685 amino acid positions. The tree was inferred with FastTree (Price et al. 2010) using the LG substitution model and gamma-distributed rates (-lg -gamma -fastest). Next, we used *treeprofiler-annotate* to annotate the resulting species phylogeny with a data table containing the average relative abundance of each taxon across 51 environments (Dmitrijeva et al. 2025). Extended taxonomic information, based on GTDB nomenclature, was also automatically inferred for all branches of the tree. Finally, *treeprofiler-plot* was used to iteratively visualize the resulting annotated tree, enabling us to explore species abundance profiles across the different habitats. Figure 3A illustrates a snapshot of this interactive exploration, where clades are color-coded by phylum, vertical colored bands delineate domain-level branches, and the right-side heatmap panel displays relative abundance profiles across various habitats. When visually navigating the tree, TreeProfiler can dynamically adjust the relative abundance heatmap for the visible branches, reflecting the average values grouped under each node (Figure 3B). At full zoom-in, the tree displays species-level resolution, showing mOTU identifiers, taxonomic labels, and original habitat-specific abundance estimates (Figure 3C).

**Figure 3.**
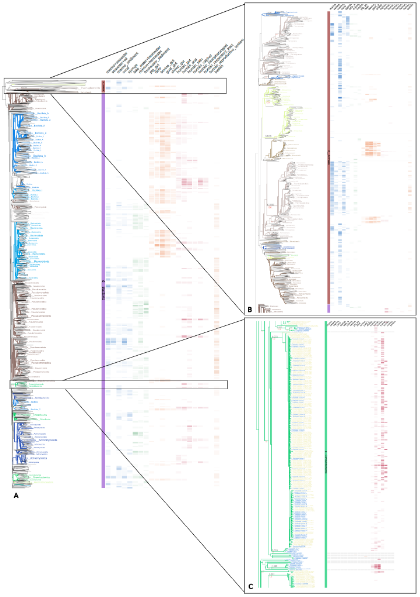
Large-scale visualization of the mOTUs reference phylogenetic tree annotated with habitat-specific abundance profiles. **A)** Overview of the full tree (124,295 leaves), with clades color-coded by phylum and a vertical band separating archaea and bacteria. The right-side heatmap shows the summarized relative abundance profiles of each branch across various habitats. **B)** Zoomed view of an archaea subtree illustrating how the heatmap layout dynamically adjusts to the visible region of the tree as you zoom, ensuring that the abundance data remains relevant and readable during interactive exploration. **C)** At full zoom, the tree provides a high-resolution view where terminal nodes representing reference species clearly display individual mOTU identifiers, detailed taxonomic labels, and precise habitat-specific abundance patterns.

## CONCLUSIONS

TreeProfiler overcomes current limitations of existing tools for phylogenetic tree profiling by offering a flexible framework for annotating, summarizing, and interactively visualizing custom metadata across large phylogenetic trees. These capabilities enable the exploration of ancestral trait distributions, lineage-specific patterns, and clade-level summaries within complex evolutionary contexts. The entire workflow is streamlined into just two command-line steps, eliminating the need for *ad hoc* pipelines and enabling the rapid analysis of large-scale datasets. Comprehensive documentation and usage examples are available at https://github.com/compgenomicslab/TreeProfiler, with code and figure reproduction instructions at https://github.com/dengzq1234/treeprofiler_paper.

## AUTHOR CONTRIBUTIONS

J.HC conceived the project. Z.D and J.HC designed the methodology. Z.D wrote the original version of the software, testing code and documentation. A.HP contributed with the domain annotation algorithm. A.A.D inferred the mOTUs reference tree and provided the mOTUs taxonomy database and abundance dataset. J.HC and Z.D wrote the manuscript, with contributions from all other authors.

## ACKNOWLEDGEMENTS

The authors want to thank Jorge Botas and Jordi Burguet-Castell for their help developing and debugging many features of the ETE Toolkit software that are used by TreeProfiler.

## FUNDING

This study was supported by FPI-Severo Ochoa predoctoral fellowship [SEV-2016-0672-18-2:PRE2018-084075] to Z.D; Research Technical Support Staff Aid [PTA2019-017593-I/AEI/10.13039/501100011033] to A.HP; The Swiss National Science Foundation through the NCCR Microbiomes (51NF40_225148) to A.A.D; CZI grant [DAF2020-218584] DOI https://doi.org/10.37921/207480vhbyqe from the Chan Zuckerberg Initiative DAF, an advised fund of Silicon Valley Community Foundation (funder DOI 10.13039/100014989); and the grant PID2021-127210NB-I00 MCIU/AEI/FEDER, UE, National Programme for Fostering Excellence in Scientific and Technical Research, to J.HC;

## SUPPLEMENTARY FIGURES

**Supplementary Figure 1.**
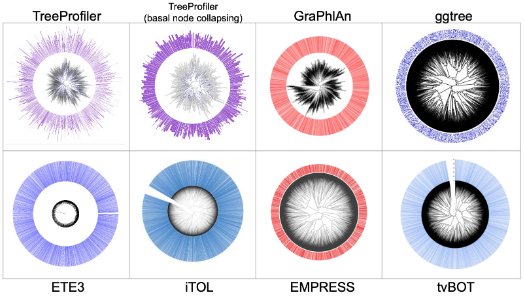
Visualization of a randomly generated tree topology with 10,000 leaves using various software tools. Each leaf has been randomly assigned a score, with half receiving a score of 0 and half receiving a score of 50. These scores are depicted by colored outer-circle bars in each visualization. At full zoom-out scale, TreeProfiler (shown in purple) displays average scores for collapsed internal branches, accurately reflecting the random distribution of values. In contrast, raw tree visualization software may misleadingly suggest that the full majority of nodes are annotated with high scores, possibly due to overlapping of score bars on collapsed nodes. Source data and program parameters used are available at https://github.com/dengzq1234/treeprofiler_paper.

## Notes

https://github.com/compgenomicslab/TreeProfiler

https://github.com/dengzq1234/treeprofiler_paper/

### Competing Interest Statement

The authors have declared no competing interest.

### Summary of Updates

We have improved the use cases with more speciflic exmaples: (i) Gene tree-based phylogenetic profiling, enabling the exploration of subfunctionalization patterns, orthologous groups, and the distribution of custom genomic features by analyzing functional diversification and domain architectures within the pyruvate flavodoxin/ferredoxin oxidoreductase gene family (13,297 genes); and (ii) Species tree-based profiling the relative abundance of 124,295 bacterial and archaeal species across 51 biomes., allowing for the interactive exploration of functional, ecological, and evolutionary patterns along species phylogenies. We also performed the comparison of peer tree visulaization softwares to showcase how treeprofiler can fill gap of limitation (Supplementary figure 1)

## REFERENCES

Asnicar F, Weingart G, Tickle TL, Huttenhower C, Segata N. 2015. Compact graphical representation of phylogenetic data and metadata with GraPhlAn. PeerJ 3:e1029.

Cantalapiedra CP, Hernández-Plaza A, Letunic I, Bork P, Huerta-Cepas J. 2021. eggNOG-mapper v2: Functional Annotation, Orthology Assignments, and Domain Prediction at the Metagenomic Scale. Mol. Biol. Evol. 38:5825–5829.

Cantrell K, Fedarko MW, Rahman G, McDonald D, Yang Y, Zaw T, Gonzalez A, Janssen S, Estaki M, Haiminen N, et al. 2021. EMPress Enables Tree-Guided, Interactive, and Exploratory Analyses of Multi-omic Data Sets. mSystems [Internet] 6. Available from: 10.1128/mSystems.01216-20

Capella-Gutiérrez S, Silla-Martínez JM, Gabaldón T. 2009. trimAl: a tool for automated alignment trimming in large-scale phylogenetic analyses. Bioinformatics 25:1972–1973.

Cromar GL, Zhao A, Xiong X, Swapna LS, Loughran N, Song H, Parkinson J. 2016. PhyloPro2.0: a database for the dynamic exploration of phylogenetically conserved proteins and their domain architectures across the Eukarya. Database 2016:baw013.

Dmitrijeva M, Ruscheweyh H-J, Feer L, Li K, Miravet-Verde S, Sintsova A, Mende DR, Zeller G, Sunagawa S. 2025. The mOTUs online database provides web-accessible genomic context to taxonomic profiling of microbial communities. Nucleic Acids Res 53:D797–D805.

Fang Y, Li M, Li X, Yang Y. 2021. GFICLEE: ultrafast tree-based phylogenetic profile method inferring gene function at the genomic-wide level. BMC Genomics 22:774.

Hernández-Plaza A, Szklarczyk D, Botas J, Cantalapiedra CP, Giner-Lamia J, Mende DR, Kirsch R, Rattei T, Letunic I, Jensen LJ, et al. 2022. eggNOG 6.0: enabling comparative genomics across 12 535 organisms. Nucleic Acids Res. 51:D389–D394.

Huerta-Cepas J, Serra F, Bork P. 2016. ETE 3: Reconstruction, Analysis, and Visualization of Phylogenomic Data. Mol. Biol. Evol. 33:1635–1638.

Ishikawa SA, Zhukova A, Iwasaki W, Gascuel O. 2019. A Fast Likelihood Method to Reconstruct and Visualize Ancestral Scenarios. Mol. Biol. Evol. 36:2069–2085.

Katoh K, Standley DM. 2013. MAFFT multiple sequence alignment software version 7: improvements in performance and usability. Mol. Biol. Evol. 30:772–780.

Letunic I, Bork P. 2024. Interactive Tree of Life (iTOL) v6: recent updates to the phylogenetic tree display and annotation tool. Nucleic Acids Res. [Internet]. Available from: 10.1093/nar/gkae268

Mendler K, Chen H, Parks DH, Lobb B, Hug LA, Doxey AC. 2019. AnnoTree: visualization and exploration of a functionally annotated microbial tree of life. Nucleic Acids Res. 47:4442–4448.

Mistry J, Chuguransky S, Williams L, Qureshi M, Salazar GA, Sonnhammer ELL, Tosatto SCE, Paladin L, Raj S, Richardson LJ, et al. 2020. Pfam: The protein families database in 2021. Nucleic Acids Res. 49:D412–D419.

Moi D, Dessimoz C. 2023. Phylogenetic profiling in eukaryotes comes of age. Proc. Natl. Acad. Sci. U. S. A. 120:e2305013120.

Parks DH, Chuvochina M, Rinke C, Mussig AJ, Chaumeil P-A, Hugenholtz P. 2021. GTDB: an ongoing census of bacterial and archaeal diversity through a phylogenetically consistent, rank normalized and complete genome-based taxonomy. Nucleic Acids Res. 50:D785–D794.

Pellegrini M, Marcotte EM, Thompson MJ, Eisenberg D, Yeates TO. 1999. Assigning protein functions by comparative genome analysis: protein phylogenetic profiles. Proc. Natl. Acad. Sci. U. S. A. 96:4285–4288.

Peng C, Guo X-L, Zhou S-D, He X-J. 2023. Backbone phylogeny and adaptive evolution of Pleurospermum s. l.: New insights from phylogenomic analyses of complete plastome data. Front. Plant Sci. 14:1148303.

Price MN, Dehal PS, Arkin AP. 2010. FastTree 2--approximately maximum-likelihood trees for large alignments. PLoS One 5:e9490.

Ravenhall M, Škunca N, Lassalle F, Dessimoz C. 2015. Inferring horizontal gene transfer. PLoS Comput. Biol. 11:e1004095.

Revell LJ. 2024. phytools 2.0: an updated R ecosystem for phylogenetic comparative methods (and other things). PeerJ 12:e16505.

Rey C, Guéguen L, Sémon M, Boussau B. 2018. Accurate Detection of Convergent Amino-Acid Evolution with PCOC. Mol. Biol. Evol. 35:2296–2306.

Ribeiro D, Borges R, Rocha AP, Antunes A. 2023. Testing phylogenetic signal with categorical traits and tree uncertainty. Bioinformatics [Internet] 39. Available from: 10.1093/bioinformatics/btad433

Rossier V, Train C, Nevers Y, Robinson-Rechavi M, Dessimoz C. 2024. Matreex: compact and interactive visualisation for scalable studies of large gene families. Genome Biol. Evol. [Internet]. Available from: 10.1093/gbe/evae100

Sayers EW, Cavanaugh M, Clark K, Ostell J, Pruitt KD, Karsch-Mizrachi I. 2018. GenBank. Nucleic Acids Res. 47:D94–D99.

Szklarczyk D, Nastou K, Koutrouli M, Kirsch R, Mehryary F, Hachilif R, Hu D, Peluso ME, Huang Q, Fang T, et al. 2025. The STRING database in 2025: protein networks with directionality of regulation. Nucleic Acids Res. 53:D730–D737.

Tran V, Ebersberger I. 2025. PhyloProfile v2 -- Exploring multi-layered phylogenetic profiles at scale. arXiv [q-bio.PE] [Internet]. Available from: 10.48550/ARXIV.2504.19710

Xie J, Chen Y, Cai G, Cai R, Hu Z, Wang H. 2023. Tree Visualization By One Table (tvBOT): a web application for visualizing, modifying and annotating phylogenetic trees. Nucleic Acids Res 51:W587–W592.

Yu G, Smith DK, Zhu H, Guan Y, Lam TT-Y. 2017. Ggtree : An r package for visualization and annotation of phylogenetic trees with their covariates and other associated data. Methods Ecol. Evol. 8:28–36.

