## Supplementary Figure 1 for "TreeProfiler: Large-scale metadata profiling along gene and species trees"

### SUPPLEMENTARY FIGURES

#### Supplementary Figure 1.

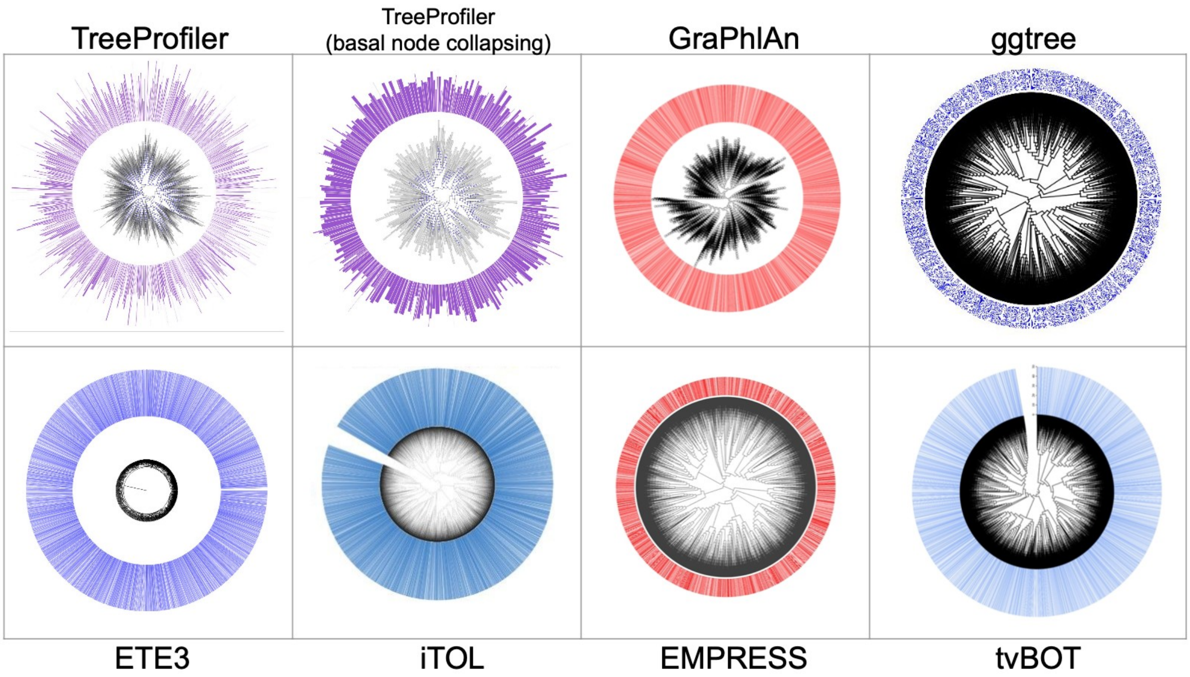


**Supplementary Figure 1**. Visualization of a randomly generated tree topology with 10,000 leaves using various software tools. Each leaf has been randomly assigned a score, with half receiving a score of 0 and half receiving a score of 50. These scores are depicted by colored outer-circle bars in each visualization. At full zoom-out scale, TreeProfiler (shown in purple) displays average scores for collapsed internal branches, accurately reflecting the random distribution of values. In contrast, raw tree visualization software may misleadingly suggest that the full majority of nodes are annotated with high scores, possibly due to overlapping of score bars on collapsed nodes. Source data and program parameters used are available at https://github.com/dengzq1234/treeprofiler_paper.
